# ADAMTS2 regulates radial neuronal migration by activating TGF-β signaling at the subplate layer of the developing neocortex

**DOI:** 10.1101/2022.08.07.502954

**Authors:** Noe Kaneko, Kumiko Hirai, Minori Oshima, Kei Yura, Mitsuharu Hattori, Nobuaki Maeda, Chiaki Ohtaka-Maruyama

## Abstract

During the development of the mammalian brain, neocortical structures are formed by the sequential radial migration of newborn excitatory neurons. The early migrating neurons exhibit a multipolar shape, but they undergo a multipolar-to-bipolar transition at the subplate (SP) layer, where extracellular matrix (ECM) components are abundantly expressed. In this study, we revealed that the TGF-β signaling-related ECM proteins, such as latent TGF-β-binding protein 1 (LTBP1) and fibrillin 2, and TGF-β receptor II (TGF-βRII) and its downstream effector, p-smad2/3, are selectively expressed at the SP layer, suggesting that TGF-β is sequestered in a latent form by forming complexes with these ECM components and then its signaling is activated by ECM remodeling. We found that the migrating multipolar neurons transiently express a disintegrin and metalloproteinase with thrombospondin motif 2 (ADAMTS2), an ECM metalloproteinase, just below the SP layer. Knockdown and knockout of Adamts2 suppressed the multipolar-to-bipolar transition of migrating neurons, and therefore, disturbed radial migration. Similar phenotypes were observed by the perturbation of TGF-β signaling in the migrating neurons. Time-lapse luminescence imaging of TGF-β signaling indicated that ADAMTS2 activates this signaling pathway in the migrating neurons during the multipolar-to-bipolar transition at the SP layer. These results suggest that the ADAMTS2 secreted by the migrating multipolar neurons activates TGF-β signaling by ECM remodeling of the SP layer, leading to the multipolar-to-bipolar transition. We propose that the SP layer plays an essential role in the radial neuronal migration as a signaling center of the developing neocortex.

**SIGNIFICANCE:** The neocortex is formed by the sequential radial migration of newborn neurons, which undergo a multipolar-to-bipolar transition at the subplate (SP) layer. The extracellular matrix (ECM) is abundantly expressed in the SP layer. However, the roles of the ECM in the SP layer have been unclear. We found that migrating neurons transiently express a disintegrin and a metalloproteinase with thrombospondin motif 2 (ADAMTS2), an ECM metalloproteinase, just below the SP layer. We show that ADAMTS2 secreted by multipolar migrating neurons activates TGF-β signaling through remodeling of the ECM in the SP layer, leading to the multipolar-to-bipolar transition. Thus, the SP layer plays an essential role in radial migration as a signaling center of the developing neocortex

## INTRODUCTION

The mammalian neocortices are densely populated with neurons of different characteristics and morphologies within their six layers, which are responsible for cognition, memory, and motor behaviors (1). Defects in the cortical layer formation processes lead to various brain malformations, and neurological and psychiatric disorders in humans (2, 3). During fetal development of the neocortex, excitatory neurons are born in the ventricular zone and travel long distances toward the pial surface, during which two migration modes, multipolar migration and locomotion, are used sequentially (4, 5) (Fig. 1A). This change of migration mode accompanies the multipolar-to-bipolar transition of migrating neurons at the subplate (SP) layer. Recently, we revealed that SP neurons (SpN), first-born, and matured neurons in the developing neocortex, form transient synapses on subsequently born multipolar neurons (MpN) (6). The glutamatergic synaptic transmission from SpNs to MpNs induces calcium signaling in the MpNs, leading to their multipolar-to-bipolar transition and change of migration mode. Although many genes have been reported to control the radial neuronal migration process (7), little is known about the molecular mechanism that induces the multipolar-to-bipolar transition after this synaptic transmission.

**Fig. 1.**
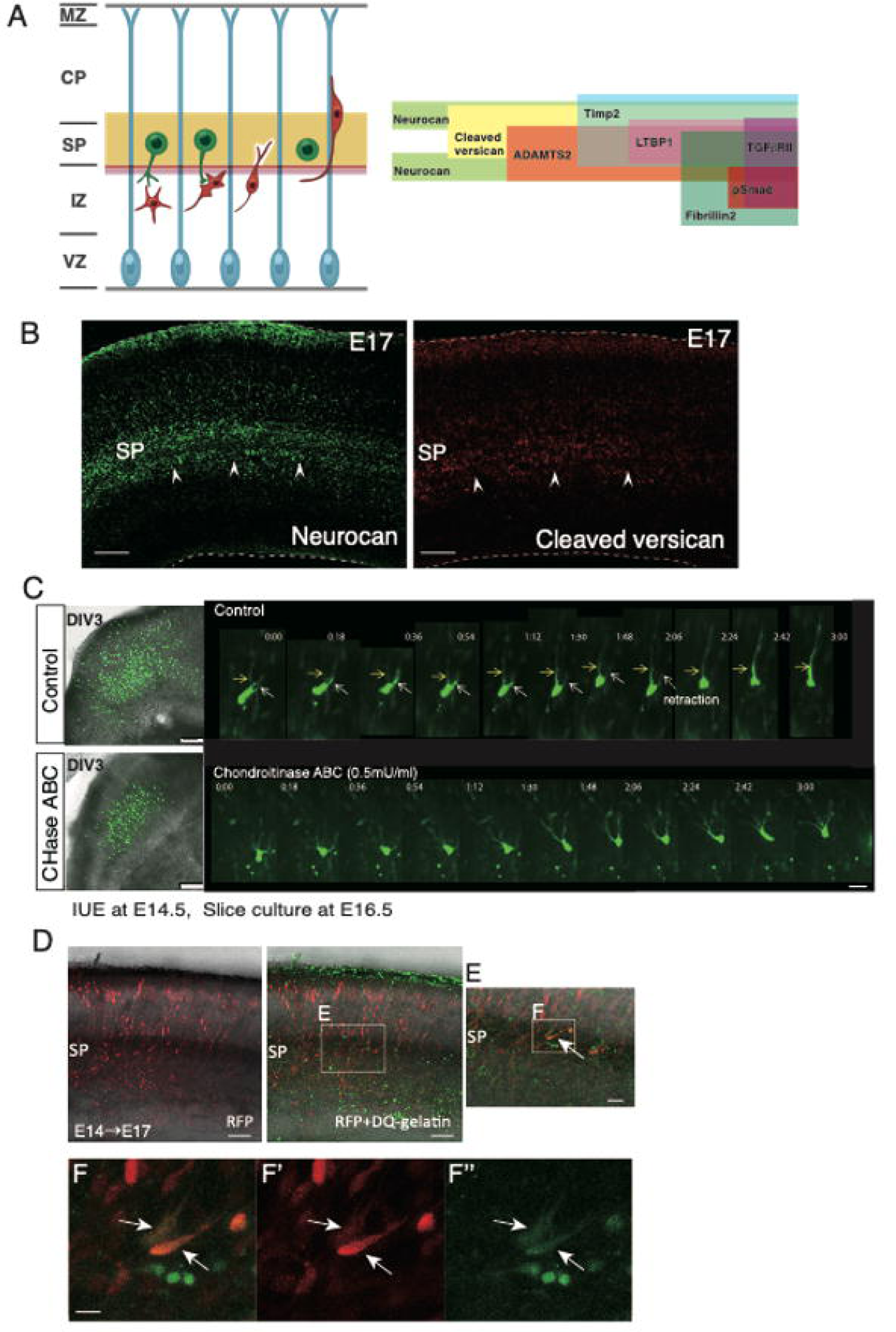
Radial neuronal migration and the ECM molecules expressed in the SP layer. **A.** A schematic model of radial neuronal migration in the neocortex (left) with a schematic representation of the distribution of various ECM molecules around the SP layer (right). **B.** Immunostaining of neurocan and cleaved versican revealed that these CSPGs were localized at the SP layer (arrow heads). **C.** CHase ABC treatment disturbed radial neuronal migration. Slices prepared from E16.5 cortices (EP of GFP at E14.5) were cultured with or without CHaseABC for 3 days (left). In the control slices, many GFP-labelled neurons migrated to the upper part of the cortex, whereas neurons were accumulated beneath the SP layer in the CHaseABC-treated slices. Time-lapse imaging of control migrating neurons revealed that a neurite became thicker when it was determined as a leading process. In contrast, CHase ABC-treated neurons remained multipolar without leading processes. **D.** *In situ* zymography using DQ-gelatin revealed that ECM protease activity (green) occurred in the SP layer. Enlarged images reveal that the migrating neurons exhibited gelatinolytic activities near the SP layer (E-F’’). Scale bars: B, D, 50 μm; C-left, 100 μm; C-right, F, 10 μm; E, 20 μm. MZ: marginal zone; CP: cortical plate; SP: subplate; IZ: intermediate zone; VZ: ventricular zone.

The SpNs reside in the SP layer, which contains abundant extracellular matrix (ECM) components, including chondroitin sulfate proteoglycans (CSPG) such as neurocan, phosphacan, and versican (8, 9). It is known that the ECM provides mechanical support to the tissues, and plays important roles in cell differentiation, adhesion, migration, and cell-cell interaction. In addition to the multidomain large glycoproteins, the ECM contains a variety of growth factors and cytokines, such as FGF and TGF-β, and regulates the stability and bioactivities of these signaling molecules (10). TGF-β is sequestered in the ECM as an inactive dimer bound to the latent TGF-β-binding protein (LTBP) by a latency-associated peptide (LAP), which is called a large latent complex (LLC). LLCs also bind to the ECM components, such as fibronectin fibers and fibrillin microfibrils, forming very large complexes (11, 12). Activation of latent TGF-β is tightly regulated by many factors, such as integrins (12) and ECM remodeling induced by proteolytic processing of LLCs (13), which leads to the release of active TGF-β. The released disulfide-bonded TGF-β dimer binds to two TGFβRII receptors, which recruits two TGFβ-RI receptors, and this heterotetrameric receptor complex initiates the downstream Smad-dependent signaling (14).

Many studies have linked the CSPGs in the SP layer with axon guidance (9, 15, 16), and our previous studies indicated that CSPGs play important roles in early neuronal polarization and migration (17, 18). However, the roles of the SP layer ECM in neuronal migration remain unclarified. As mentioned above, the SP layer is a strategically important region where the multipolar-to-bipolar transition occurs during radial neuronal migration. Thus, we investigated how the ECM of the SP layer is involved in the morphological changes of migrating neurons. In this study, we found that the expression of ADAMTS2 (a disintegrin and metalloproteinase with thrombospondin motif 2) in the migrating neurons was significantly up-regulated just below the SP layer during the multipolar-to-bipolar transition and is required for the radial neuronal migration. Although ADAMTS2 is well-known as a procollagen-cleaving enzyme and a causative gene for Ehlers-Danlos syndrome (EDS), a skin disorder caused by abnormal collagen fibrillogenesis (19, 20), Bekhouche et al. recently revealed that TGF-β signaling is the primary target of ADAMTS2 (21). The substrates of ADAMTS2 included LTBP-1, TGF-βRIII, and ADAMTS2 activated TGF-β signaling in the human fibroblasts. Therefore, we focused on TGF-β signaling as a downstream candidate of ADAMTS2. We found that TGF-β signaling is transiently activated in the migrating neurons just below the SP layer, in which ADAMTS2 is essential for the activation of the signaling. We suggest that the ECM remodeling induced by ADAMTS2 secreted by migrating neurons transiently activates TGF-β signaling at the SP layer, which leads to the multipolar-to-bipolar transition and the change of the migration mode (Fig. S6).

## RESULTS

### ECM molecules and remodeling in the SP layer

Firstly, we examined the distribution of ECM molecules in the developing neocortex. Immunohistochemistry and *in situ* hybridization analyses revealed that CSPGs, including neurocan, cleaved versican, and phosphacan/PTPζ (PTPRZ1), are highly expressed at the SP layer (Fig. 1B, Fig. S1A). In addition, fibronectin 1, collagen XIα1, LTBP 1, and fibrillin 2 are also expressed at the SP layer (Figs. S1A and B).

The ECM is not a static structure but is dynamically remodeled during various biological processes, especially during development (22–24). To examine whether ECM remodeling is involved in radial neuronal migration, we cultured cortical slices of mice from embryonic day 16 (E16), in which neurons were electroporated with GFP-expression plasmids at E14, for 3 days in the presence of chondroitinase ABC (CHase ABC). CHase ABC is a bacterial enzyme that degrades chondroitin sulfate and hyaluronan, and thus, breaks down ECM structures, which may mimic the ECM remodeling. The results showed that there was a significant delay in neuronal migration when CHase ABC was added (Fig. 1C, left panels). Time-lapse imaging showed that the multipolar-to-bipolar transition hardly occurred, and the leading processes were not easily determined in the treated neurons (Fig. 1C, right panels, Movie 1). These results suggested that the normally regulated ECM plays an essential role in the multipolar-to-bipolar transition and radial neuronal migration.

Next, we tried to visualize the ECM proteolytic activity in the developing cortex by using *in situ* zymography with DQ-gelatin. DQ gelatin is a fluorogenic substrate consisting of highly quenched, fluorescein-labeled gelatin (25). Upon proteolytic degradation, green fluorescence is revealed, and thus, can be used to localize proteolytic activities in the ECM. As a result, we found that the migrating neurons exhibited gelatinolytic activities in the SP layer, which were suppressed by the metalloproteinase inhibitors, 1,10-phenanthroline and GM6001 (Fig. 1D, Fig. S2). These results indicated that the ECM components and proteolytic activities are present in the SP layer, supporting the hypothesis that the ECM remodeling contributes to the multipolar-to-bipolar transition during radial neuronal migration.

### Adamts*2* is expressed in radially migrating neurons

Next, we used microarray data to investigate which genes are involved in the ECM remodeling. To date, we have performed microarray analyses to reveal the changes in gene expression of the migrating neurons during radial migration (6). We introduced GFP-expression plasmid into the neural progenitor cells at E14 by *in utero* electroporation, and then the GFP-labelled neurons were separated by fluorescence-activated cell sorting (FACS) analysis at E15, E16, and E17. In this scheme, MpNs, neurons undergoing multipolar-to-bipolar transition, and bipolar neurons can be collected at E15, E16, and E17, respectively (Fig. 2A). As a result of the microarray analyses of these neurons, 2,377 probes with significant changes in the expression levels were extracted (6). We re-analyzed these data, and extracted genes that have ECM-related GO (Gene Ontology) terms. The expression levels of many ECM-related genes changed significantly during radial neuronal migration (Fig. S3A).

**Fig. 2.**
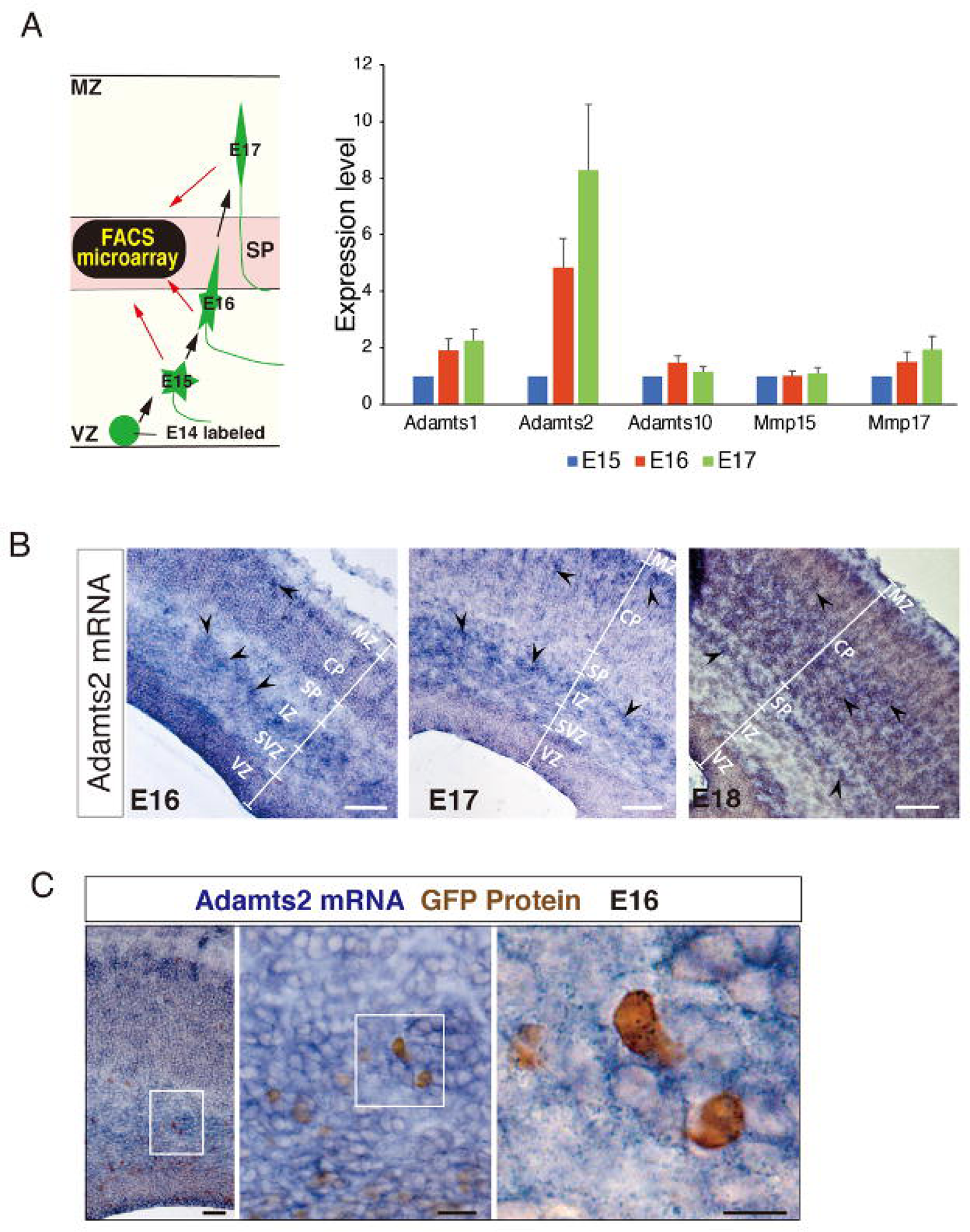
Migrating neurons express ADAMTS2 in the SP layer. **A.** Left: A schematic representation of the FACS microarray analyses of migrating neurons. Right: FACS microarray analyses revealed that the expression of Adamts2 mRNA was increased during radial migration**. B.** Adamts2 mRNA was localized at the upper part of IZ, the lower part of the SP layer, and the CP. **C.** Double staining of ADAMTS2-ISH and anti-GFP antibody-IMH revealed that GFP-positive migrating neurons express *Adamts2* mRNA. Scale bars: 100μm for B, 50μm for C-left, 20μm for C-middle, 10μm for C-right.

Among these genes, we focused on the ECM metalloproteinases as enzymes that remodel ECM structures and identified ADAMTS2 as a gene whose expression is markedly increased during radial migration. It was apparent that ADAMTS2 expression elevated more rapidly during the multipolar-to-bipolar transition than the other metalloproteinases such as ADAMTS1, ADAMTS10, MMP15, and MMP17 (Fig. 2A). ADAMTS2, ADAMTS3, and ADAMTS14 form procollagen N-proteinase subfamilies (21, 26, 27), and therefore, we compared their expression levels by quantitative PCR (Fig. S3B). We found that the expression of ADAMTS2 was markedly up-regulated during migration among the subfamily members.

*In situ* hybridization experiments indicated that ADAMTS2 is expressed in the SP layer (Fig. 2B). To confirm that migrating neurons express ADAMTS2, we performed double staining of the cortical sections, in which migrating neurons were labeled with GFP at E14 by *in utero* electroporation and fixed at E16. As a result, the expression of *Adamts2* mRNA (blue) was confirmed in the GFP-positive migrating MpNs (brown) around the bottom part of the SP layer (Fig. 2C). Therefore, we decided to further analyze the function of ADAMTS2 in the regulation of neuronal migration.

### ADAMTS2 regulates neuronal migration

To investigate the involvement of ADAMTS2 in radial neuronal migration, we performed knockdown and overexpression of ADAMTS2 in migrating neurons. First, we examined the knockdown efficiency by si-RNA (*Silencer* Select predesigned siRNA for *Adamts2*; s103675, ambion) using NIH3T3 cells. As a result, 10 μg/μl of si-RNA was the most efficient in suppressing the Adamts2 mRNA expression (Fig. S4A). We introduced si-RNA against Adamts2 into the neural progenitor cells along with GFP-expression plasmids by *in utero* electroporation at E14, and the brains were fixed at E17 and E18. We quantified the distribution of GFP-positive neurons in the cortices and found that Adamts2 knockdown suppressed the radial migration, and many neurons lingered below the SP layer (Fig. 3A). These migration defects were rescued by introducing the Adamts2-expression plasmid along with siRNA (Fig. 3B).

**Fig. 3.**
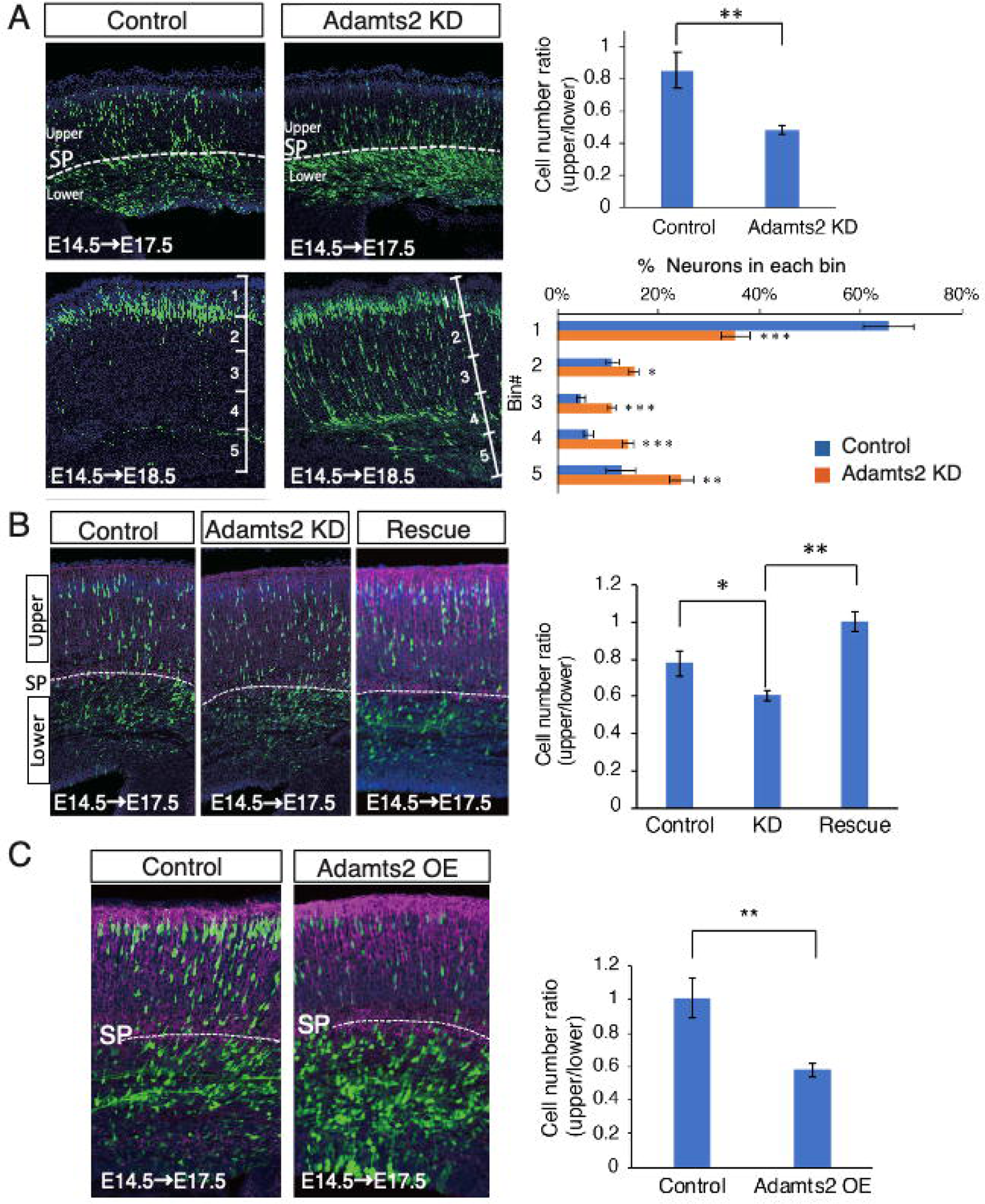
*Adamts2* is involved in the regulation of radial neuronal migration. **A**. GFP expression plasmids were co-electroporated with si-RNA for *Adamts2* at E14.5, and the distribution of GFP-positive neurons was analyzed at E17.5 and E18.5. Compared with the control, knockdown of *Adamts2* resulted in the retardation of radial neuronal migration. Many neurons were stacked just below the SP layer at E17.5 and E18.5. A part of the neurons remained in the middle of migration at E18.5 (N = 18). **B.** The impairment of migration by *Adamts2* knockdown was rescued by co-electroporation of *Adamts2* expression vectors (N = 5). **C.** Neuronal migration was impaired by overexpression of *Adamts2* (N = 7).

Interestingly, Adamts2 overexpression also impaired migration (Fig. 3C). Then, we further examined Adamts2 KO mice (BJ6/ADAMTS2 Δ28KO mice) (28). In the heterozygous Adamts2 KO mice, most migratory neurons were stagnant below the SP layer, which was similar to the phenotype of Adamts2 knockdown (Fig. S4D). These results suggested that an appropriate level of Adamts2 expression is required for radial neuronal migration. However, migration defects were not observed in the homozygous Adamts2 KO mice (Fig. S4D). This may be explained by the genetic compensation in the homozygous mice that lack the Adamts2 gene.

To observe the details of the migration defect, LifeAct-GFP plasmid, which can label F-actin, was introduced into the migrating neurons along with RFP plasmid by *in utero* electroporation at E14. After two days, the brains were dissected, and the cortical slices were cultured for time-lapse observation (Figs. S4B and C, Movie 2). In contrast to the control, where many migrating neurons changed from the multipolar to bipolar shape, many migrating neurons in Adamts2 knockdown remained multipolar and showed altered F-actin dynamics. In the control slices, F-actin was detected in the multiple processes of MpNs, but after the multipolar-to-bipolar transition, F-actin was concentrated in the leading processes of bipolar neurons undergoing locomotion. On the other hand, in Adamts2 knockdown neurons, the dynamics of F-actin remained the same as in the control MpNs for at least 24 hours, and F-actin did not assemble in the leading processes (arrow heads in Fig. S4C, Movie 2). These results indicated that ADAMTS2 is involved in the multipolar-to-bipolar transition of migrating neurons.

### TGF-β signaling-related proteins are localized at the SP layer

Recently, it was demonstrated that TGF-β signaling-related proteins, such as LTBP1 and TGF-βRIII, are potential substrates of ADAMTS2, suggesting the participation of this protease in the control of TGF-β activity (21). It was also proposed that the polarity transition of newborn neurons mechanically resembles the epithelial-mesenchymal transition (EMT) that occurs during cancer cell invasion, in which TGF-β signaling is deeply involved (29). Therefore, we focused on TGF-β signaling as a candidate target for ADAMTS2.

First, we examined the localization of TGF-β signaling-related proteins in the developing neocortex: phospho-Smad2/Smad3 (p-smad2/3), a major downstream effector of TGF-β signaling, tissue inhibitor of metalloproteinase 2 (Timp2), an inhibitory factor of TGF-β signaling, and TGF-βRII. Immunohistochemistry revealed that p-smad2/3 and Timp2 were localized in the lower and the upper parts of the SP layers, respectively (Figs. 4A-H). Immunoreactivity to TGF-βRII was also located in the SP layer (Fig. 4I,I’).

**Fig. 4.**
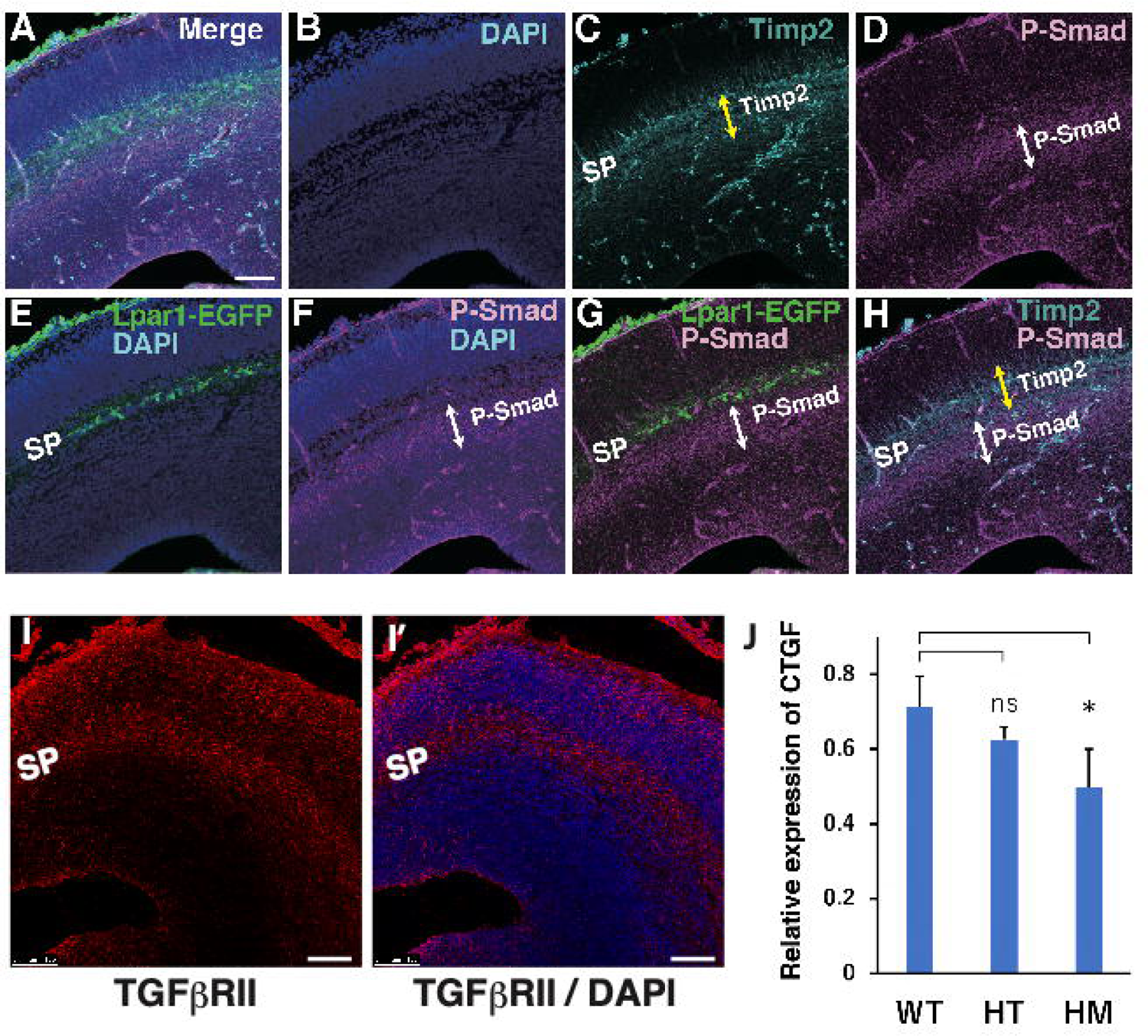
TGF-β signaling-related proteins are localized at the SP layer. **A-H.** Cortical sections from Lpar1-EGFP mice, in which SP neurons expressed EGFP, were immunostained with antibodies against TGF-β signaling-related proteins. Timp2 and p-Smad were expressed at the upper part and the lower part of the SP layer, respectively. **I, I’.** TGF-βRII immunoreactivities were localized at the SP layer and the upper part of the intermediate zone. **J.** The expression of CTGF, a direct downstream target of TGF-β signaling, was down-regulated in the cerebral cortex of Adamts2 KO mice. The expression levels of CTGF were measured by Q-PCR using mRNAs isolated from the cerebral cortex. All sections were E15.5.

Next, we examined the expression of CTGF in the cerebral cortices of Adamts2 KO mice by quantitative PCR. CTGF is a direct downstream target gene of TGF-β signaling (30–32) and is expressed selectively in the SP layer (33, 34). The results showed that the expression of CTGF in the cerebral cortex was significantly reduced in homozygous KO mice compared with that of wild-type mice (Fig. 4J), suggesting that TGF-β signaling was down-regulated in the brains of Adamts2 KO mice.

### Effects of TGF-β signaling on neuronal migration

To investigate the effects of TGF-β signaling on radial neuronal migration, TGF-βRII was overexpressed by *in utero* electroporation at E14, and the brains were fixed at E17. The results showed that migrating neurons stagnated below the SP layer when TGF-βRII was overexpressed (Fig. 5A). Yi et al. also reported that knockout of TGF-βRII in migrating neurons resulted in a delayed migration and axonal loss phenotype (35). We then analyzed the effects of the inhibitors of TGF-β signaling on the actin dynamics and migration of neurons. We prepared slice cultures of the cerebral cortex at E16 after introducing RFP and LifeAct-GFP expression plasmids by *in utero* electroporation at E14. Then, TGF-βRI inhibitors (RepSox and LDN-212854) were added to the culture medium, and the movements of migrating neurons were monitored by time-lapse imaging for 3 days. In the presence of inhibitors, the neurons remained multipolar and stagnated under the SP layer, whereas many control neurons migrated across the SP layer and exhibited a bipolar shape (Figs. 5B and C). Furthermore, inhibitor-treated neurons showed abnormal dynamics of F-actin, and the formation of leading processes was impaired (Movie 3), the phenotype of which was similar to that of Adamts2 knockdown neurons (Fig. S4C, Movie 2). These results suggest that TGF-β signaling transiently activated around the SP layer is involved in the multipolar-to-bipolar transition of migrating neurons. In the inhibitor-containing slices, the axon elongation of migrating neurons was also significantly impaired, suggesting that TGF-β signaling also regulates axon determination and elongation of newborn neurons (Fig. S5) (35).

**Fig. 5.**
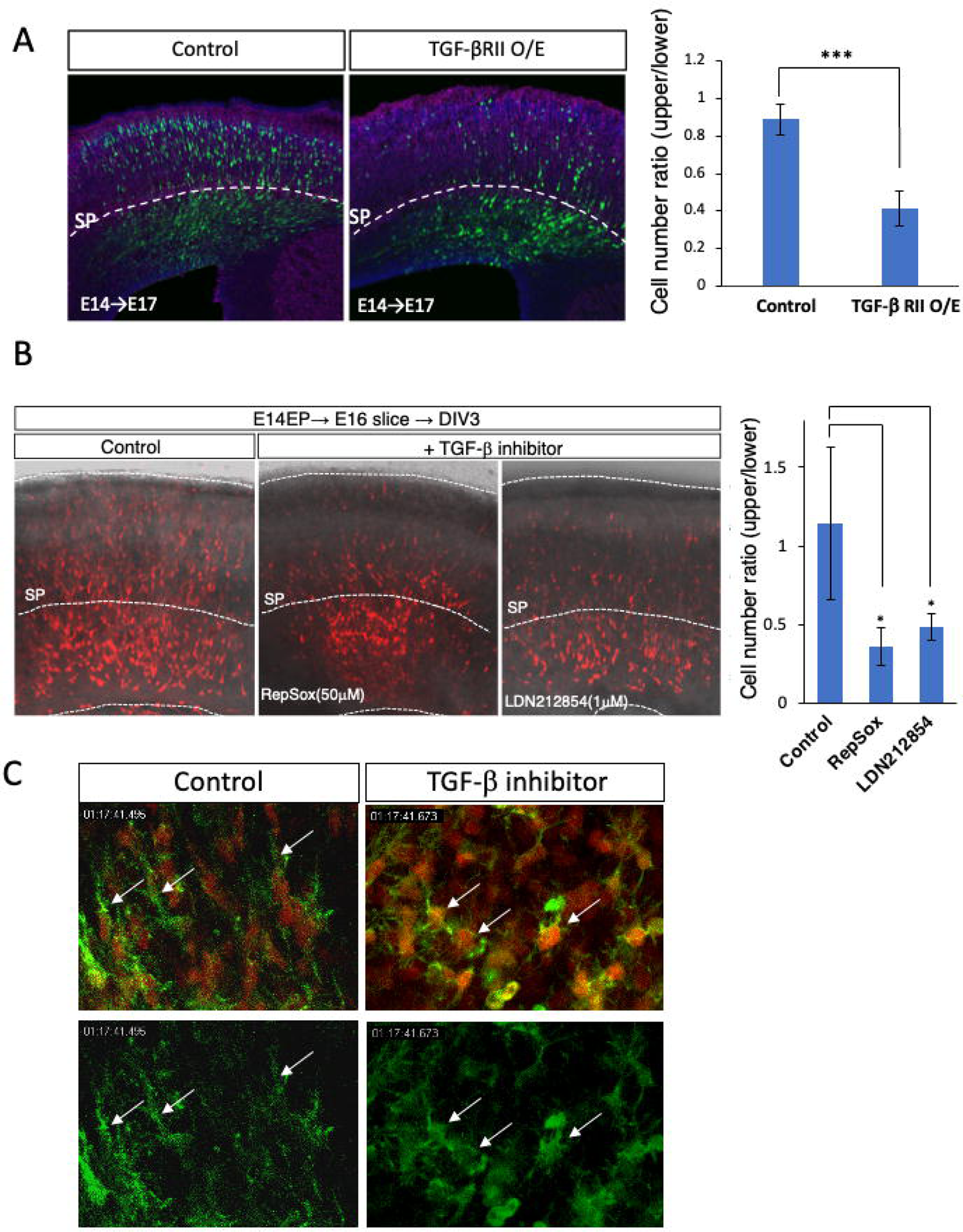
Perturbation of TGF-β signaling impaired radial neuronal migration. **A.** Overexpression of TGF-βRII in the migrating neurons impaired radial neuronal migration. **B.** Inhibitors of TGF-β receptor (50 μM RepSox and 1 μM LDN212854) disturbed radial neuronal migration in the cultured slices. **C.** Selected images from the time-lapse recordings of F-actin dynamics shown in Movie 3. CAG-LifeAct and RFP plasmids were electroporated at E14.5. The brain slices were prepared at E16.5 and cultured in the presence or absence of 50 μM RepSox. The multipolar-to-bipolar transition was impaired in the presence of the inhibitor.

### ADAMTS2 turns on the TGF-β signaling at the SP layer and leads to morphological changes in migrating neurons

We visualized TGF-β signaling by luminescence imaging to confirm that TGF-β signaling is turned on in the migrating neurons at the SP layer. We created a construct in which two DNA sequences that are reactive to p-smad2/3 (TGF-β-reactive elements) were placed in tandem, upstream of the Emerald-Luc coding sequence that is fused with the PEST sequence, a protein-destabilizing signal (Fig. 6A). In the cells transfected with this construct, luciferase is only expressed after activation of TGF-β signaling. The plasmid was introduced into the cortical neurons by *in utero* electroporation, and the brain slices were observed by luminescence imaging. The luminescence signals were detected at the lower part of the SP layer and the upper part of the intermediate zone, and the signals were suppressed in the presence of RepSox, indicating that they were specific (Figs. 6B-D, G, Movie 4). The luminescence signals were significantly reduced in the Adamts2 knockdown neurons, indicating that ADAMTS2 is required for the activation of TGF-β signaling (Figs. 6E and G, Movie 4). The enlarged time-lapse images demonstrated that the MpNs in the lower part of the SP layer transiently showed strong luminescence signals, which disappeared after their multipolar-to-bipolar transition (Fig. 6F). These results suggest that ADAMTS2 activates TGF-β signaling in the MpNs around the bottom part of the SP layer, which leads to the multipolar-to-bipolar transition.

**Fig. 6.**
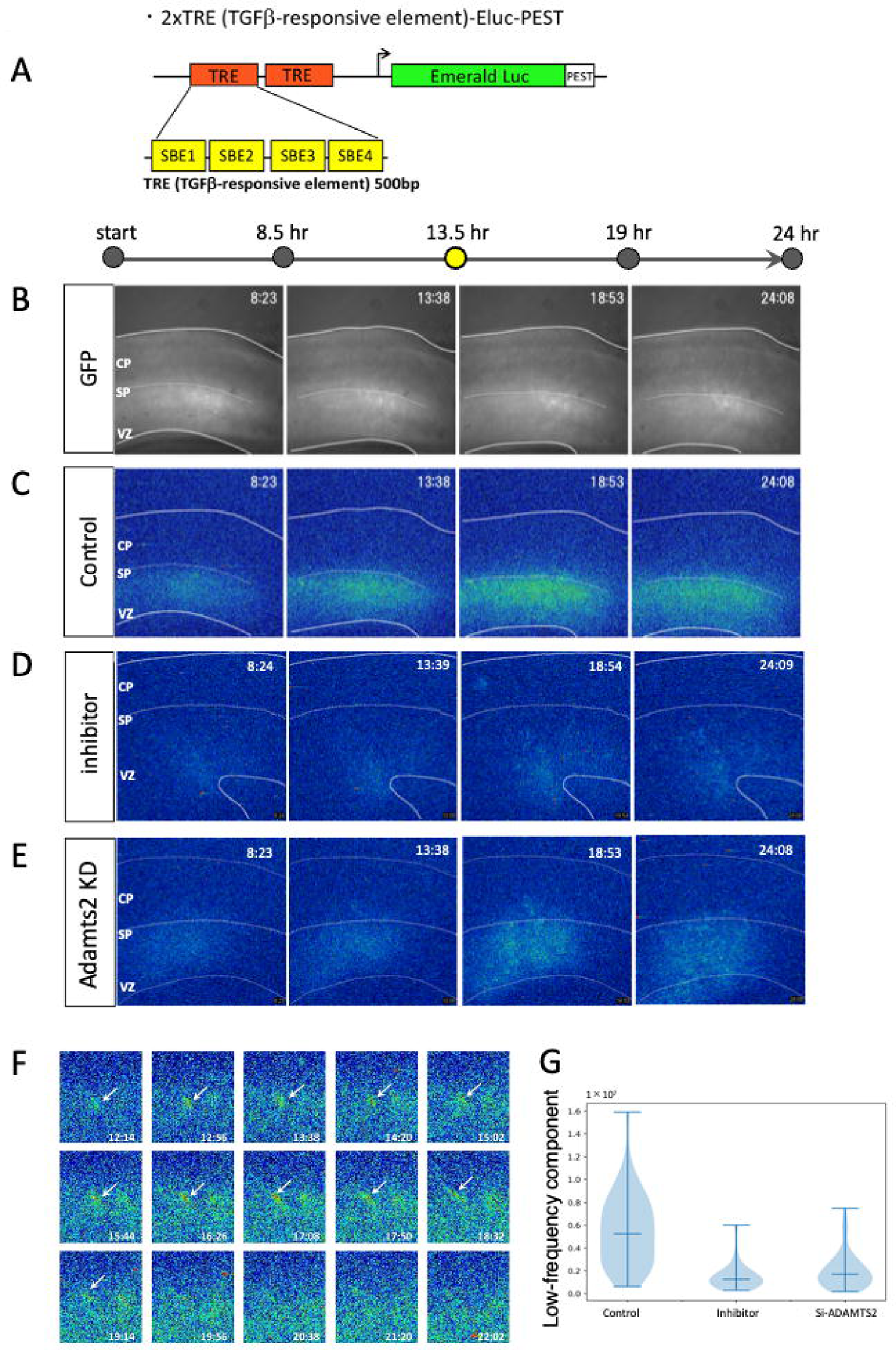
Time-lapse imaging of TGF-β signaling during radial neuronal migration. **A.** The plasmid construct used for luminescent imaging of TGF-β signaling. TGF-β-responsive elements were tandemly inserted into the upstream of Emerald Luc conjugated with the PEST sequence. **B-E.** 2xTRE-Eluc-Pest plasmids were electroporated at E14.5 along with GFP expression plasmids. The luminescence imaging was performed using cultured slices prepared at E16.5. Compared to the control (C), the luminescence signals were diminished when 50 μM RepSox was added (D), or Adamts2 si-RNA was co-electroporated (E). **F**. Enlarged images revealed that a control migrating neuron transiently showed strong luminescence emission during the multipolar-to-bipolar transition from 12 to 18 hours. **G**. Quantification of the luminescence signals. The data were taken 4.5 hours after the start of imaging.

## DISCUSSION

In this study, we revealed that the SP layer is enriched with many ECM components and works as a strategically important region where TGF-β signaling is activated. ADAMTS2 that is secreted by the migrating MpNs activates TGF-β signaling in these neurons, which induces multipolar-to-bipolar transitions and locomotion. Unlike many signaling molecules, TGF-β is sequestered in the ECM in a latent form, and it should be activated by multiple steps before binding to the receptors (11–14). The secretion and extracellular regulation of TGF-β is considered to proceed as follows. After translation, pro-TGF-β dimerizes and is then bound to LTBP by a disulfide bond in the endoplasmic reticulum. In the *trans*-Golgi network, the pro-TGF-β dimer is cleaved into a TGF-β dimer and LAP to form the small latent complex (SLC), in which LAP is non-covalently bound to TGF-β. The LLC is then secreted and sequestered in the ECM by binding to fibronectin fibers and fibrillin microfibrils until it is released by activators. Activation and release of TGF-β from the latent complex can be mediated by numerous factors such as proteinases, integrins, and some chemicals. In the SP layer, LTBP1, fibrillin 2, fibronectin, neurocan, and versican are richly expressed (Fig. 1A). The C-terminal G3 domain of versican interacts with fibronectin and fibrillin 2, and the N-terminal G1 domain interacts with hyaluronan (36). Neurocan also interacts with hyaluronan (37). Thus, it is likely that hyaluronan, versican, neurocan, fibrillin 2, and fibronectin form a macromolecular complex in the SP layer where LLC is sequestered. Based on this consideration, we propose the following hypothetical model for the ADAMTS2-mediated activation of TGF-β signaling (Fig. S6). Around the bottom part of the SP layer, the migrating MpNs transiently secrete ADAMTS2, which cleaves TGF-β-related ECM proteins such as LTBP1 and versican. After these initial cleavages, active TGF-β is released from the ECM, which initiates the activation process of TGF-β signaling, leading to the multipolar-to-bipolar transition and switching of the migration mode (Fig. S6). At present, we do not know the detailed structure of the ECM in the SP layer. However, CHase ABC treatment disturbed the multipolar-to-bipolar transition at the SP layer (Fig. 1C), suggesting that CSPGs and hyaluronan are the critical components of the SP layer ECM. Future studies are necessary to reveal the ultrastructure of the SP layer ECM, including whether fibrillin-2 and fibronectin are assembled into fiber structures. We also do not know the activation processes of TGF-β after ECM cleavage by ADAMTS2. One possibility is that ADAMTS2 loosens the SP layer ECM, which enables the migrating neurons to access TGF-β. Alternatively, ECM cleavage by ADAMTS2 may trigger multi-step processing of LLCs, which leads to the release of active TGF-β dimer. However, the step of TGF-β activation following ECM cleavage by ADAMTS2 remains unclarified.

A previous study reported that no apparent defects were observed in the layer formation and structural maintenance of the adult neocortex of Adamts2 KO mice (28). We also observed a few defects in the radial neuronal migration process in the cortices of homozygous Adamts2 KO mice. However, the defects of multipolar-to-bipolar transitions and radial migration were observed in the cortices of heterozygous Adamts2 KO mice. Similar defects were observed in the Adamts2 knockdown neurons, and even in the Adamts2 overexpression neurons. These findings suggest that strictly regulated expression of Adamts2 is necessary to control TGF-β signaling and radial migration. Under Adamts2 knockdown and heterozygous Adamts2 KO conditions, MpNs may not be able to secrete enough ADAMTS2 to release sufficient amounts of TGF-β due to deficient ECM cleavage in the SP layer. On the contrary, Adamts2 overexpression may cause excessive cleavage of the ECM, leading to the overactivation of TGF-β signaling. Overexpression of TGF-βRII resulted in migration defects similar to the Adamts2 overexpression, suggesting that excess TGF-β signaling disturbs radial migration (Fig. 5A). It is also known that disturbances of TGF-β signaling is associated with Loeys-Dietz syndrome, in which aortic aneurysms are associated with arterial tortuosity (38). Thus, the activities of TGF-β signaling in the SP layer are strictly controlled by the transient expression of an appropriate level of ADAMTS2 by MpNs. On the other hand, in the Adamts2 homozygous KO mice, the absence of this enzyme might be compensated for by the expression of the other metalloproteinases. ADAMTS2 is a member of the 19-gene ADAMTS family and is evolutionarily and structurally most closely related to ADAMTS3 and 14. ADAMTS2, 3, and 14 recognize and cleave similar sites on their substrates and enhance each other’s cleavage activity (21). Therefore, ADAMTS3 and ADAMTS14 may have rescued the migration defects of Adamts2 homozygous KO mice. Triple knockout of ADAMTS2, 3, and 14 may be required for further functional studies.

It is well known that ADAMTS2, 3, and 14 cleave the N-terminal propeptide of fibrillar procollagen. However, in addition to having fragile skin due to the abnormal collagen structures, Adamts2 KO homozygous mice have been genetically linked to male infertility, reduced numbers of female pups, and childhood strokes, suggesting a sizeable unknown substrate repertoire beyond procollagen N-terminal cleavage (28). In fact, secretome analyses of human fibroblasts using N-terminal amine isotope labeling of the substrate revealed many candidate substrates for ADAMTS2, 3, and 14 (21). These include LTBP1, CTGF, fibronectin, and versican, which are richly expressed in the SP layer. Similar approaches may reveal other novel substrates for ADAMTS2 in the SP layer. Furthermore, ADAMTS2 and 3 cleave Reelin at the specific site (N-t-cleavage) and abolish its biological activity (28, 39). These findings show that ADAMTS2 has a wide range of functions in the developing cortex.

It is apparent that the activation of TGF-β signaling is a multi-step process that requires numerous factors such as proteinases and protease inhibitors. The immunoreactivity to p-smad2/3 was confined to the bottom part of the SP layer and the upper part of the intermediate zone (Fig. 4D). Zymography experiments also indicated that gelatinolytic activities were concentrated around the bottom part of the SP layer (Fig. 1D). These findings suggest that the activities of ECM proteinases that activate TGF-β signaling are spatially controlled in a strict manner. The immunoreactivity to Timp2 was detected at the upper part of the SP layer (Fig. 4C), and thus, this protease inhibitor may inhibit ECM proteinases, including ADAMTS2, and suppress the excess TGF-β signaling activity. Spatial and temporal regulation of many ECM components may contribute to the regulation of radial neuronal migration (Fig. 1A).

It was previously reported that TGF-β signaling forms a concentration gradient in the ventricular zone of the developing cerebral cortex and regulates early axon specification events of MpNs (35). We also observed that some of the TGF-β signaling-related molecules, such as LTBP1and ADAMTS2, are commonly expressed in both the SP layer and the ventricular zone (Fig. 2B, Fig. S1B). On the other hand, the immunoreactivity to p-smad2/3 was low in the ventricular zone compared with that in the SP layer (Fig. 4D). This is consistent with the report showing that early axon specification utilizes a non-smad Par6-dependent TGF-β signaling pathway (35). Thus, migrating neurons utilize the non-canonical and canonical TGF-β signaling pathways in the different cortical layers to regulate the early axon specification and multipolar-to-bipolar transition, respectively. Accordingly, our results of the perturbation experiments regarding TGF-β signaling may include the defects that occurred early in the ventricular zone. To exclude this possibility in future experiments, we will need to use an expression plasmid with the NeuroD promoter, which drives the expression only in the differentiated neurons.

This study shows that the ADAMTS2 and TGF-β signaling function near the bottom part of the SP layer, and are involved in the change of morphology of migrating neurons from the multipolar to bipolar shape. The SP layer is unique to mammals and plays a critical role in the developmental formation of the neocortex (40, 41). Our finding of the SP layer as a signaling center for the developing neocortex may aid the elucidation of multiple mechanisms underlying the evolution of the neocortex. Furthermore, failure of radial neuronal migration causes various diseases such as brain dysplasia and psychiatric disorders (2, 7, 42), and the results of this study will contribute to understanding the pathogenesis of these diseases.

## Acknowledgements

We thank Kuniko Kohyama (Child Brain Project) for her advice on *in situ* zymography with DQ-gelatin, Song Xianghe (Neural Development Project) for analyzing data, and Aiko Odajima for her technical support in image processing. We thank members of the Neural Development Project of Tokyo Metropol. Inst. Med. Sci. for discussion and comments on the study. This work was supported in part by the JSPS KAKENHI-Grants (17K07428, 19H04795, 20H03270 to C.O.-M., and 16K07077 to N. M.), and AMED under Grant Number JP21gm1310012, Y2018 Research Grant from Takeda Science Foundation, The Naito Foundation, FY2020 Research grant from the Novartis Foundation, Brain Science Foundation, FY2021 Research Grant from Yamada Science Foundation, KOSE Cosmetology Research Foundation, The Mitsubishi Foundation, Astellas Foundation for Research on Metabolic Disorders to C.O.-M. The microarray data are available in Gene Expression Omnibus (GEO) database under accession number GSE102911.

## Materials and Methods

### Experimental animals

All animals were treated in accordance with the Tokyo Metropolitan Institute of Medical Science Animals Care and Use Committee guidelines. Pregnant ICR mice were purchased from Japan SLC and used for *in utero* electroporation and microarray analyses. Lpar1-EGFP mice (Tg(Lpar1-EGFP)GX193Gsat) were obtained from MMRRC. ADAMTS2 KO mice (BJ6/ADAMTS2 Δ28) that contain a deletion of 28 bp downstream of the start codon of exon 1 were generated using the CRISPR/Cas9 system as previously reported (23).

### Antibodies

The primary antibodies used for immunostaining were mouse anti-neurocan (43), rabbit anti-cleaved versican (ab19345, Abcam), rabbit anti-Fibrillin-2(bs12166R, Bioss), rabbit anti-LTBP1 (ab78294, Abcam), chicken anti-GFP (ab13970, Abcam), rabbit anti-MAP2 (AB5622, Merck Millipore), mouse anti-TIMP2 (ab1828, Abcam), rabbit anti-pSmad3(ab52903, Abcam), and rabbit anti-TGF-βRII (bs0117R, Bioss).

The secondary antibodies used were Alexa Fluor 488-conjugated donkey anti-chicken IgY (IgG) (703-545-155, Jackson ImmunoResearch), Alexa Fluor 546-conjugated donkey anti-rabbit IgG (A10040, Themo Fisher Scientific), and Cy5-conjugated donkey anti-mouse IgG (715-175-150, Jackson ImmunoResearch). For double staining of *in situ* hybridization (*Adamts2* mRNA) and immunohistochemistry (GFP), biotin-conjugated goat anti-chicken IgY (ab97133, Abcam) and streptavidin-conjugated HRP (SA-5004, Vector) were used. The antibodies were used at a 1:500 dilution unless otherwise noted.

### Plasmid construction

All the cDNA fragments described below were cloned into the EcoRI sites of the pCAG-GS expression vector. pCAG-tRFP was generated by cloning the turboRFP (Evrogen) coding sequence into pCAG-GS. For the construction of pCAG-Adamts2 and pCAG-TGFβR2, the Adamts2 or TGFβR2 coding sequence was amplified by PCR using mouse *Adamts2* and *TGFβR2* ORF plasmids (pCMV-*Adamts2*, Harvard plasmid clones; pCMV-*TGFβR2*, Addgene) as templates. Amplified cDNA fragments were inserted into the EcoRI sites of pCAG-GS plasmid. pCAGGS-EGFP was a gift from Dr. Ayano Kawaguchi. Si-RNA-resistant plasmid for *Adamts2*(CAG-mr-*Adamts2*) was constructed as follows. AccIII-NcoI fragments of the Adamts2 coding sequence that contains the si-RNA target site were cut out and replaced by synthetic DNA fragments with inserted codon mutations that do not change the amino acids. pCAG-Lifeact plasmid was constructed from pEGFP-C1Lifeact-EGFP (Addgene). NheI-BglII fragments of the coding region were subcloned into the CAG-GS vector. TGFβ signal monitoring plasmid, p2XTRE-Eluc-PEST, was constructed with the pEluc(PEST)-test (TOYOBO). Four TGFβ-responsive elements (SBE) are inserted in tandem, upstream of the Emerald-Luc coding sequence.

### In *utero* electroporation

The pregnant mice were deeply anesthetized with sodium pentobarbital at 50 mg/kg, and the uterine horns were exposed. A plasmid DNA solution (3-5 μg/μl) in HEPES buffered saline, pH 7.2 (HBS) containing 0.01% Fast Green, was injected into the lateral ventricle with a glass micropipette using a microinjector IM-31 (Narishige). Approximately 1-2 μl of the plasmid solutions were injected into E14.5 brains. The heads of E14.5 embryos in the uterus were placed between a tweezer-type electrode, 5 mm in diameter (LF650P5, BEX), and then five electric pulses (35 V, 50 ms in duration at intervals of 950 ms) were delivered using a CUY21E electroporator (BEX). After electroporation, the uterine horns were returned to the abdominal cavity to allow the embryos to continue development.

### Immunohistochemical staining

The embryonic brains were dissected and fixed in 4% paraformaldehyde (PFA)/ PBS overnight at 4°C. The tissues were cryoprotected in 15% sucrose/ PBS for 2-3 hr, followed by 30% sucrose/ PBS overnight at 4°C. The brains were then embedded in OCT compound (Tissue Tek) and cut into 20-μm-thick sections using a cryostat HYRAX C50 (Zeiss). The sections were soaked in PBS for 5 min and pre-incubated with 0.01% Triton X-100/PBS for 15 min, which was then incubated overnight at 4°C with primary antibodies diluted with PBS containing 0.5% skim milk. After washing three times with PBS, the sections were incubated with species-specific anti-IgG antibodies conjugated to Alexa Fluor 488, Alexa Fluor 546, or Cy5. Then, sections were mounted with PermaFluor (Thermo Scientific) after DAPI staining (5 μg/ml, Sigma-Aldrich). Images were captured using the Zeiss LSM710, LSM780, and Leica SP8 confocal microscopes.

### Slice cultures

Embryonic brains electroporated with various expression constructs were dissected at E15.5 or E16.5, and embedded in 3% low-melting agarose gels prepared in HBS. Embedded brains were cut into 300-μm-thick coronal slices using a vibratome VT1200S (Leica). The slices were placed on the insert membrane (PICMORG50, Merck Millipore) and then incubated in Neurobasal medium (Gibco) supplemented with B27 (Gibco) and antibiotics (Antibiotic-Antimycotic, Gibco) under 5% CO_2_ and 60% O_2_.

### Chondroitinase ABC treatment

The stock solution, ChondroitinaseABC (10 U/ml) (Seikagaku Corporation), was first diluted 10-fold and then 1/2000 of it was added to the medium to a final concentration of 0.5 mU/ml in the culture medium (Neurobasal supplemented with B-27).

### *In situ* zymography

The EnzCheckTM Gelatinase/Collagenase Assay Kit (Thermo Fisher E12055) was used for *in situ* zymography. DQ-gelatin was diluted to a 100mg/ml final concentration with the reaction buffer (0.05 M Tris-HCl, 0.15 M NaCl, 5 mM CaCl 2, 0.2 mM sodium azide, pH 7.6). For the preparation of cortical culture slices, RFP-expressing plasmids were electroporated *in utero* at E14, and culture slices were prepared at E17. Time-lapse imaging was performed after incubating the prepared slices with DQ-gelatin solution for 30 min. 1,10-phenanthroline (Sigma131377) and GM6001 (SelleckS7157) were added to a final concentration of 10 mM and 50 μm, respectively.

### Time-lapse imaging

The slices were cultured using stage top incubators Chamlide TC (Live Cell Instrument) for SP5 and STXG-GSI2X (TOKAI HIT) for SP8 under 5% CO_2_ and 60% O_2_. Time-lapse recordings were performed using a Leica SP5 or SP8 inverted confocal microscope with a 20× long-operation objective lens (HC PL FLUOTAR, L 20x/0.40 CORR, Leica). The maximum intensity projection was generated from 10-15 Z-stack images with 10-μm intervals at each time point.

### Si-RNA mediated knockdown

Silencer Select Pre-designed si-RNA(Amnion) for Adamts2 (s103675) and negative control (AM4635) were used for knockdown experiments. Si-RNAs were introduced to embryonic ventricles of the cerebral cortex by *in utero* electroporation along with GFP-expressing plasmids.

### *In situ* hybridization

RNA probe for *Adamts2* mRNA was prepared with the 1.3kb BamH1-EcoNI fragment of the *Adamts2* coding sequence. The fragment was subcloned into pBluescriptKS+ and riboprobe was transcribed with digoxigenin-labeled dUTP. *In situ* hybridization and double staining with *in situ* hybridization and immunohistochemistry were performed as previously described (44). The primary antibody used in this study was anti-GFP (1:500) (ab13970, Abcam).

### Q-PCR

Q-PCR was performed as described previously (44), where 2 μg of total RNA was used to make 1^st^ strand cDNA. The mRNA levels were quantified by real-time PCR using the ABI 7500 real-time PCR system.

### TGFβ inhibitors

The TGFβ inhibitors used in this study were RepSox (E616452, Selleck) and LDN-212854 (S7147, Selleck).

### Luminescent imaging of TGFβ signaling

The Live Cell Bioluminescence Imaging System Cellgraph AB-3000B (ATTO) was used for bioluminescence imaging. TGF-β signaling monitor plasmid, 2xTRE-Eluc-Pest, was electroporated into the mouse cortex at E14 and cultured slices were prepared at E16. D-Luciferin potassium salt (Fuji film) was added to the culture medium at the final concentration of 5 mM.

### Quantification of luminance values

The analysis was performed using python 3.8.0. The luminescence image was analyzed in a grid of 16 x 14 squares, and the average luminance value of each square was produced. The average of the signals along the time axis for each grid was Fourier transformed. We used these values to visualize changes in luminance over time. A low-pass filter was performed by adding frequencies of 30∼75 Hz for each grid to remove high outliers due to cosmic rays. The location of the SP layer was then confirmed in the brightfield image, and luminance values were obtained for only the top and bottom four rows of the SP layer. Because of the high background immediately after the start, the first 4.5 hours were not used for the analysis. The frequency domain was then summed from 2∼30 Hz for each grid, and the results were represented in a violin plot.

### Statistical analysis

Statistical analyses were performed using Graph Pad Prism 6.0. All data were expressed as the mean ± SEM, and Student’s *t*-tests (unpaired two-tailed) were used to compare the means of the two groups.

---

**Fig.S1.**
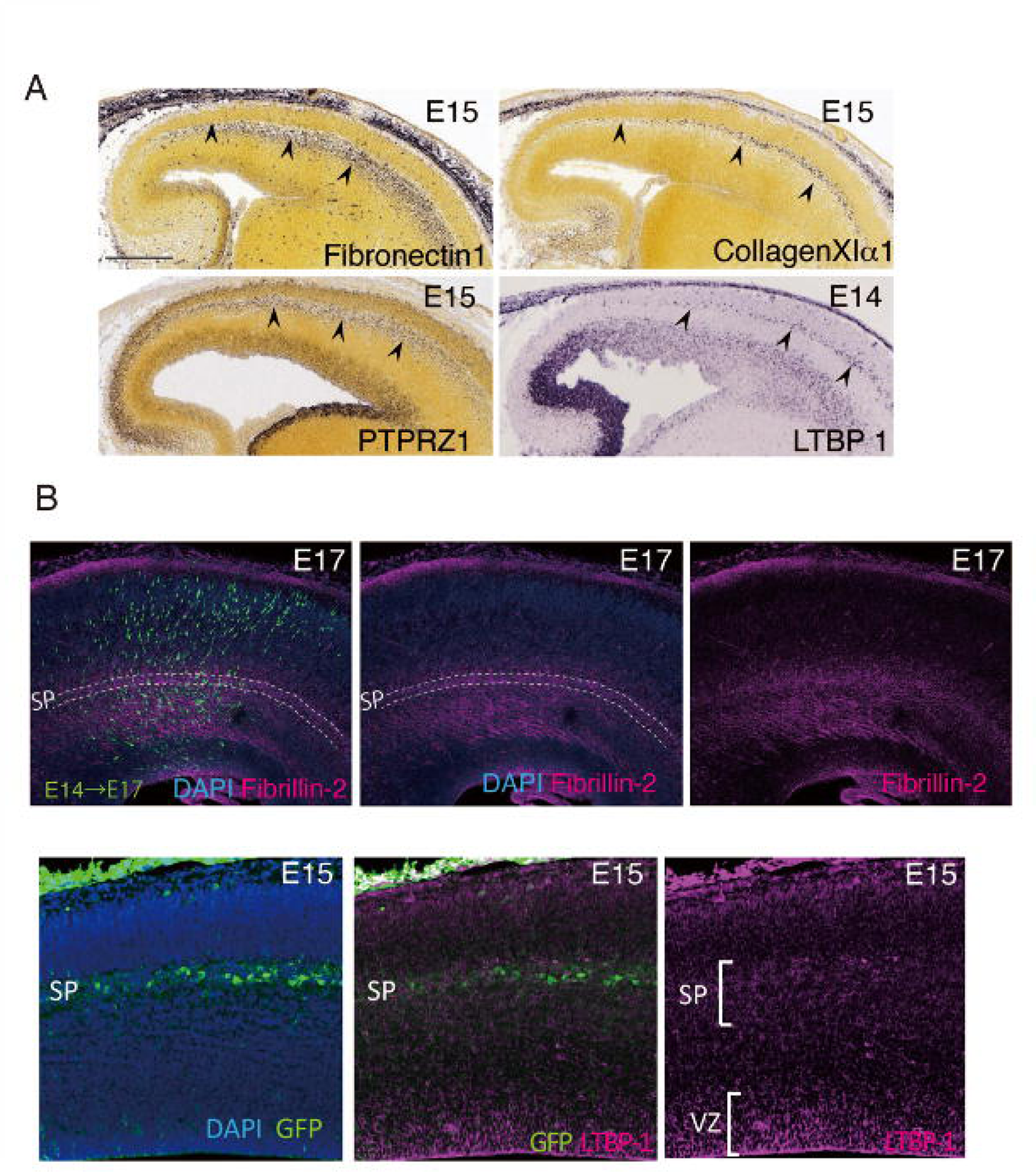
The subplate layer is rich in ECM component. A. *In situ* hybridization databases revealed that mRNAs of genes encoding ECM proteins are localized at the subplate layer in the developing mouse cortex. The data for Fibronectin 1, Collagen Xia and PTPRZl are from Allen brain atlas and the data for LTBPl is from Gene Paint. B. lmmunohistochemistry of Fibrillin-2 and LTBP-1 revealed that these ECM proteins are expressed abundantly in the subplate layer and its vicinity.

**Fig.S2.**
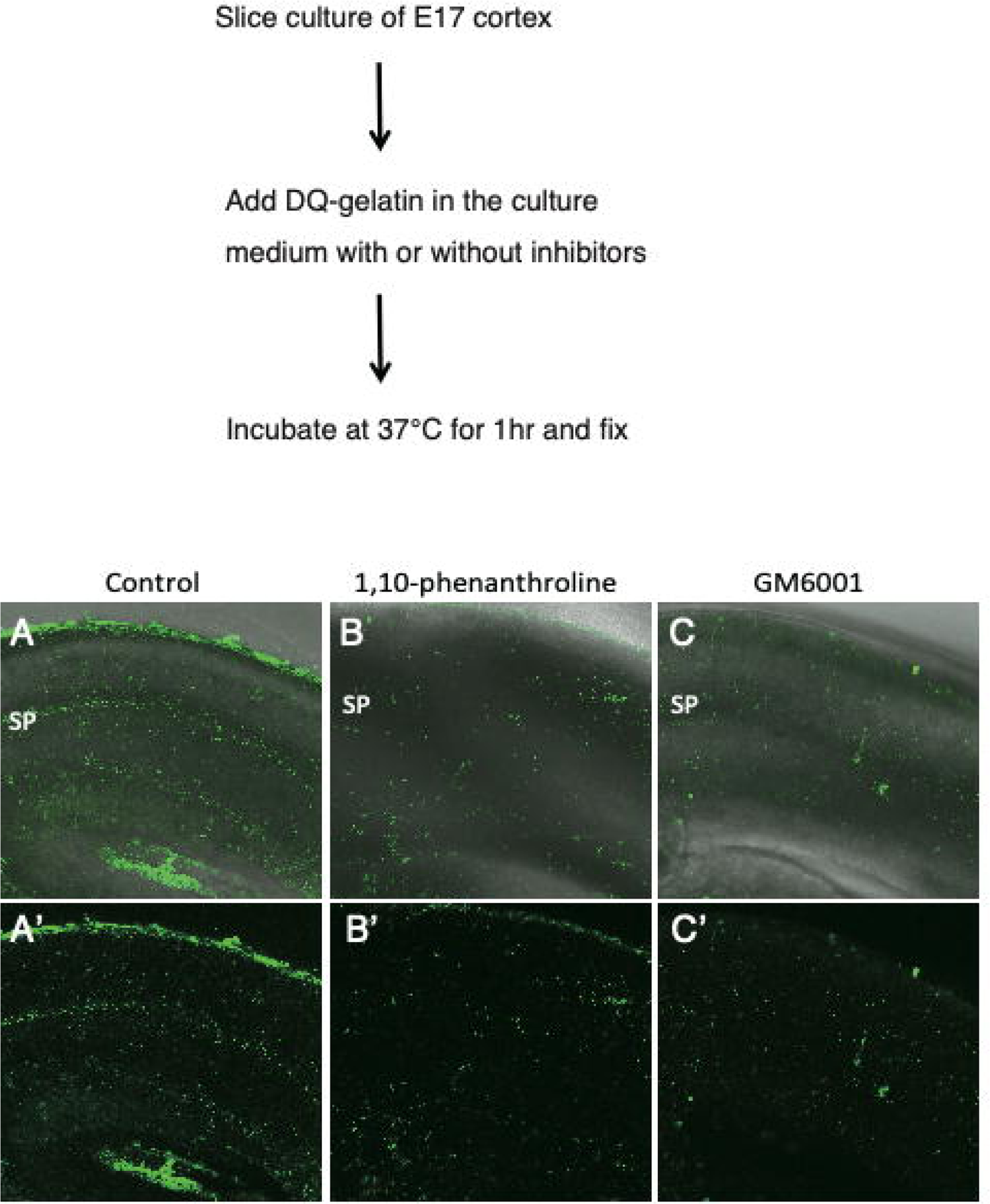
*In situ* zymography using DQ-gelatin revealed ECM protease activities In the SP layer. **A,** FITCsignals that were visualized by proteolysis of DQ-gelatin. The MZ and the SP layer are distinctly FITC-positive (A,A’). **8-C’,** The slices incubated with metalloproteinase inhibitors (1,10-phenanthroline and GM6001) did not show these signals.

**Fig.S3.**
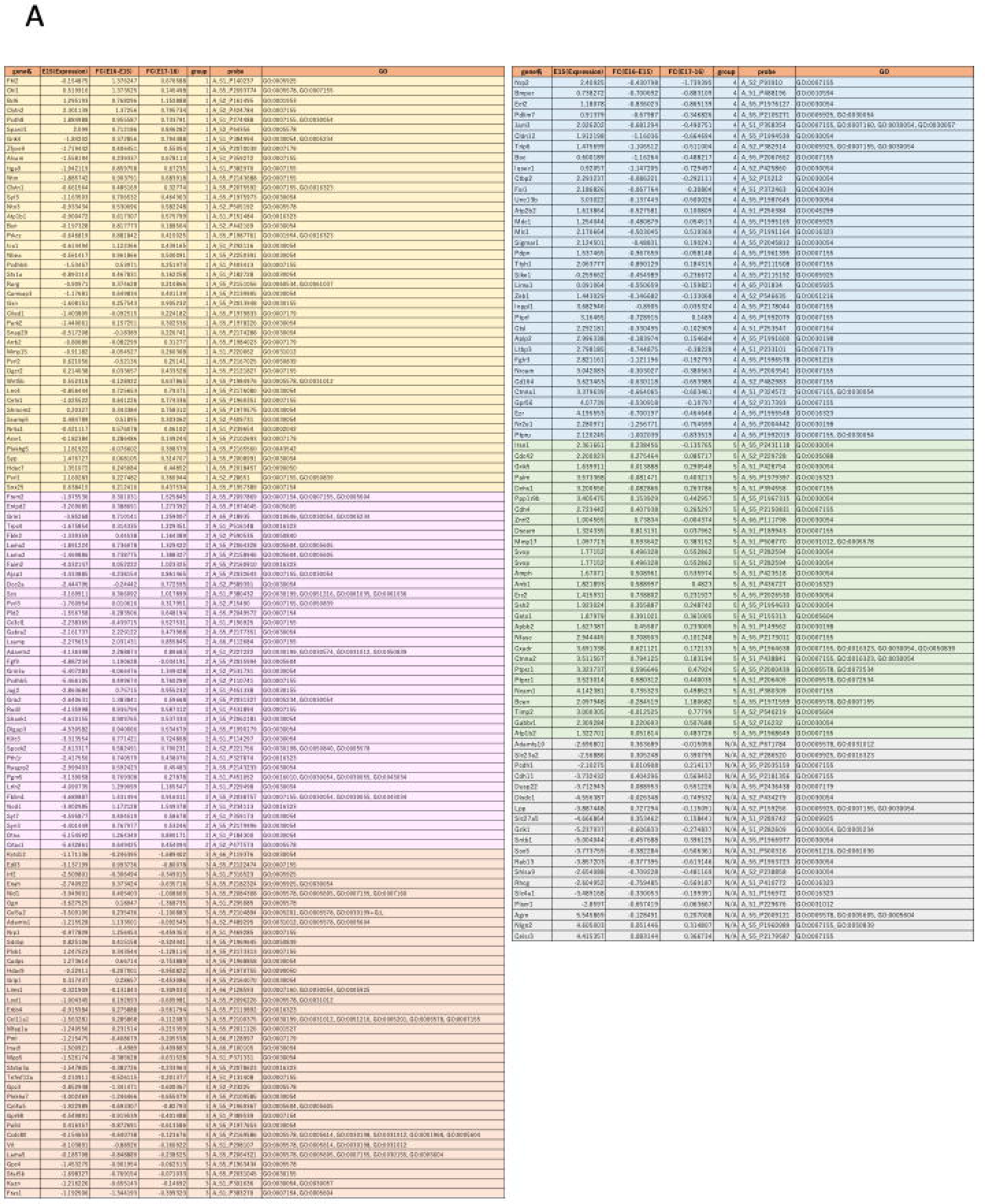
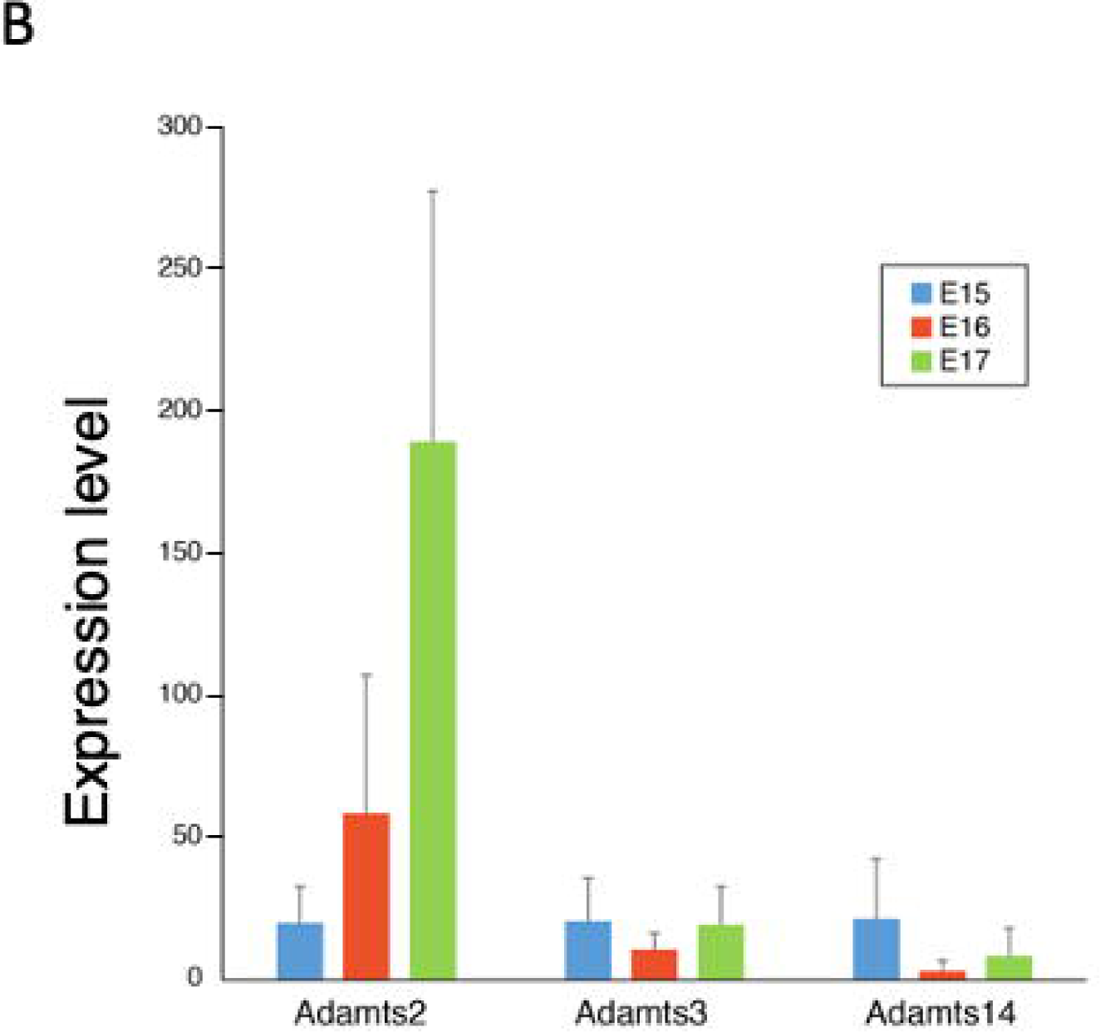
ECM-related genes, whose expression changed during radial neuron migration. **A.** Genes with GO terms related to ECM, whose expressions changed during migration, were extracted from the microarray data of migrating neurons (6). The gene list was divided into five groups. **B.** A comparison of the expression levels of ADAMTSZ, 3 and 14 subfamilies in FACS-sorted migrating Neurons (N=3). In particular, the expression level of ADAMTSZ was found to be significantly upregulated during migration.

**Fig. S4.**
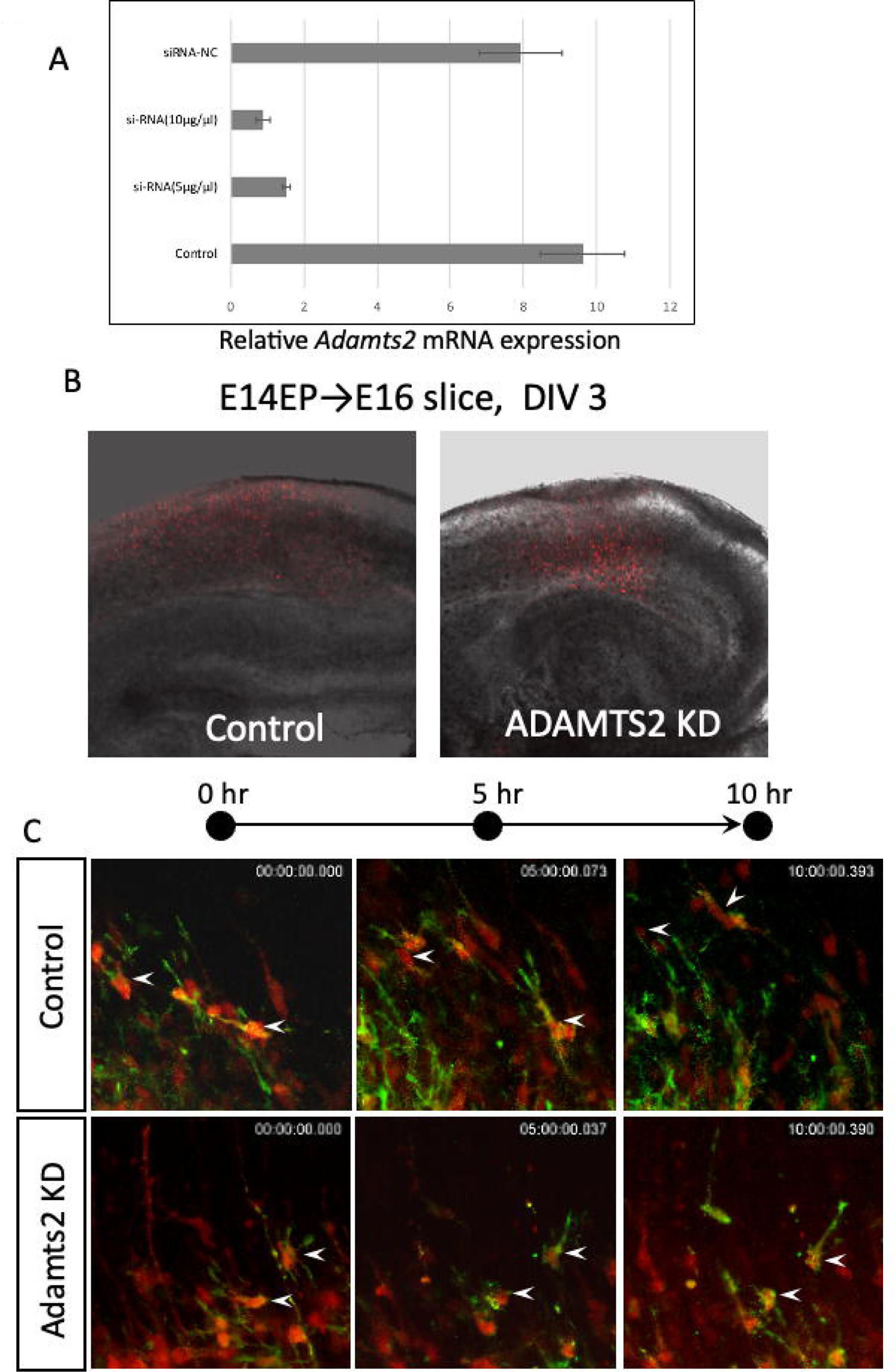
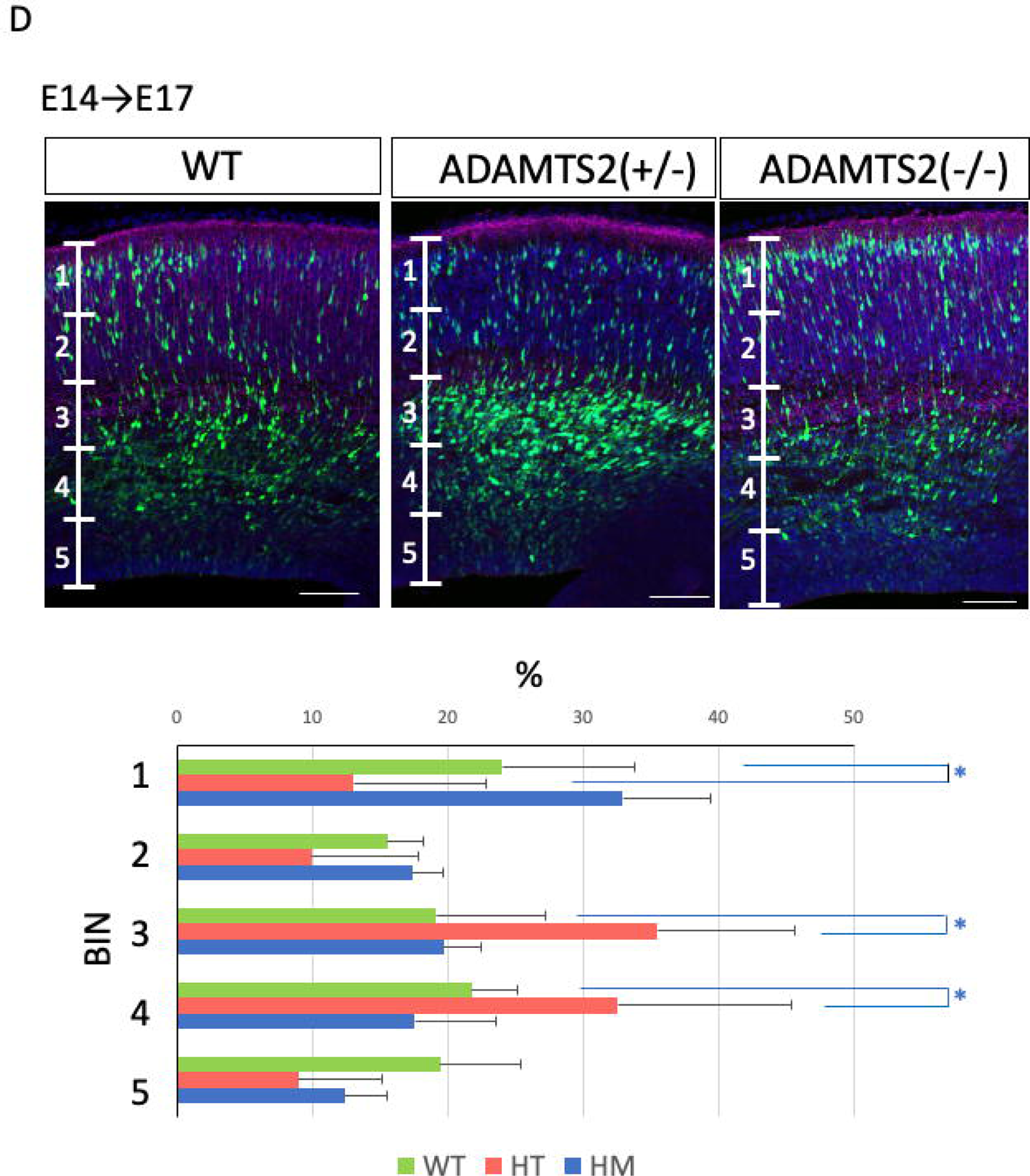
Time-lapse imaging revealed that *Adamts2* knockdown neurons were Impaired in multipolar-blpolar conversion. **A.** Confirmation of si-RNA knockdown efficiency and their optimal concentration for Adamts2 using cultured cells. B.CAG-lifeAct(F-actin labeling) and RFP plasmids were electroporated at E14.5 and cultured slices were prepared at E16.5. Time-lapse imaging was performed for 10hrs. Images of cultured slices after 3days in culture. Many cells were remained under the SP layer as multipolar morphology. C. The dynamics of F-Actin was disturbed (movie 2). **D.** Adamts2 KO mice showed impaired migration of neurons. The heterozygous (HT) mice had a more significant migration defects than the homozygous(HM) mice. N=6 for WT and HT, N=4 for HM Scale bars; 100µm

**Fig. S5.**
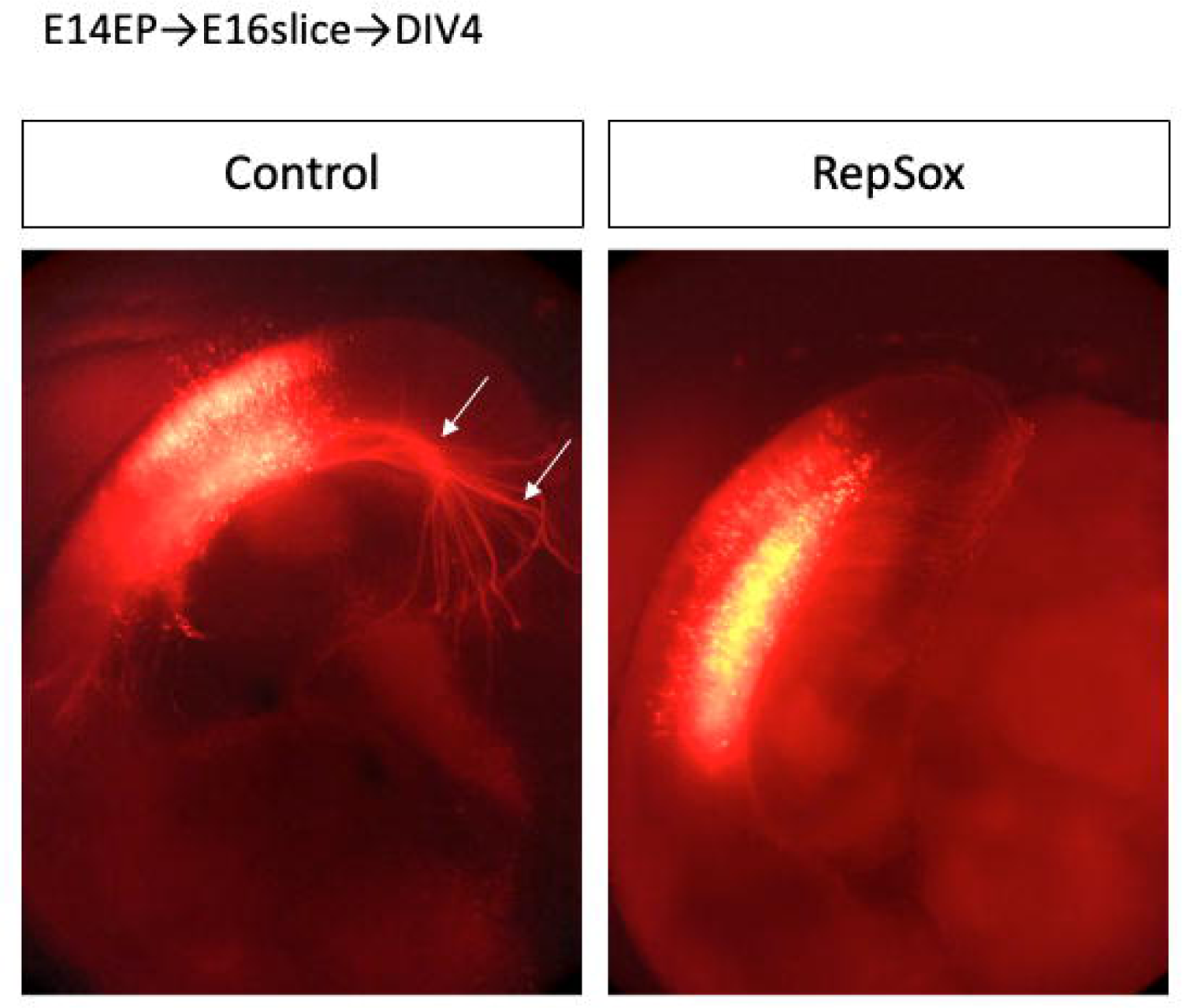
TGFβR inhibitor impaired migration and inhibited axon elongation. RFP expression plasmids were introduced into neural progenitor cells by in utero electroporationat E14, and cultured slices were prepared at E16. After time-lapse imaging, slices were fixed at DIV4.

**Fig. S6.**
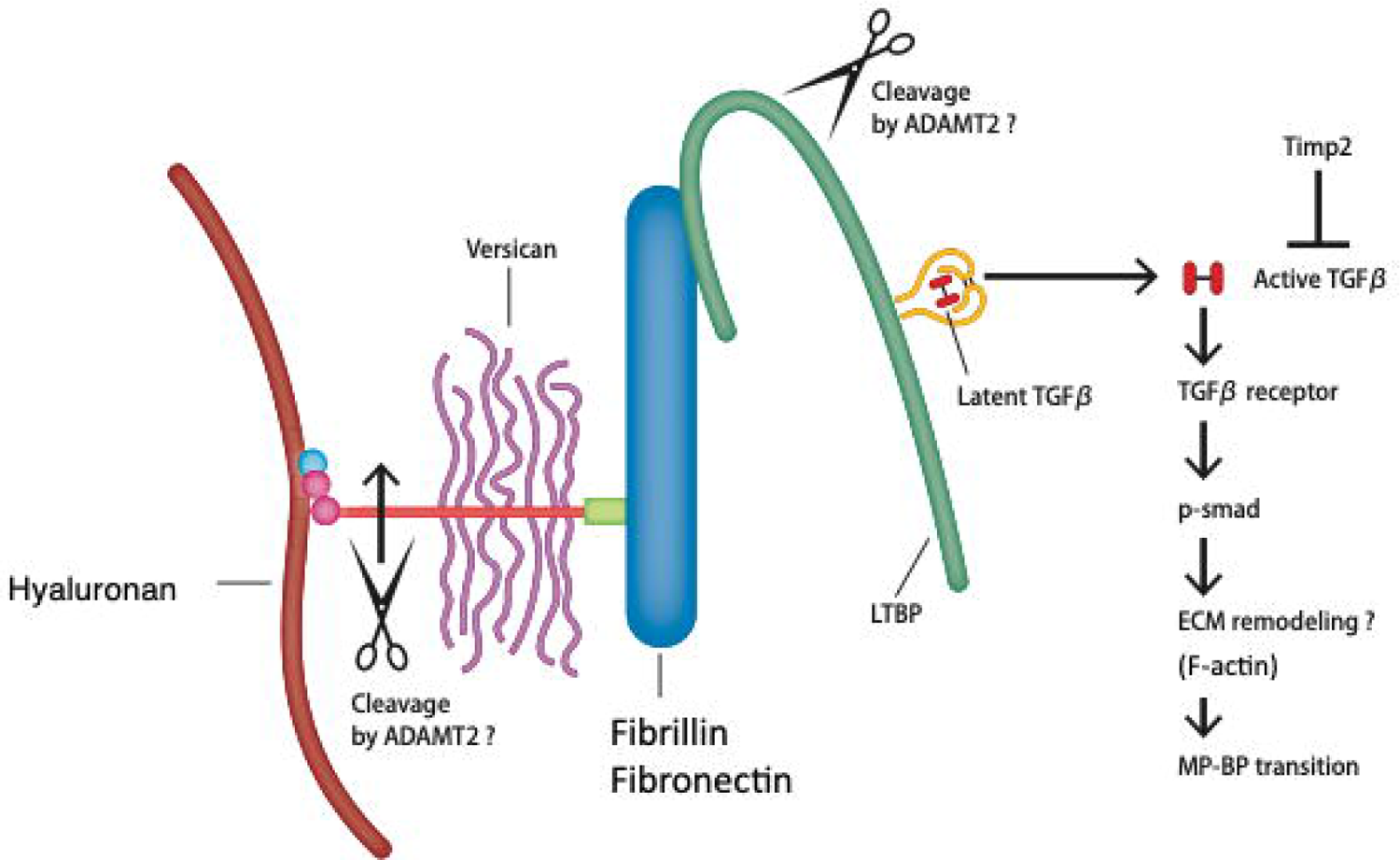
hypothetical model. The migrating multipoltransiently secrete ADAMTS2 around the bottom part of thar neurons e SP layer, which cleaves TGF-β-related ECM proteins such as LTBPl and versican. After these initial cleavages, active TGF-β-is released from the ECM, which initiates the activation process of TGF-β signaling, leading to the multipolar-to-bipolar transition and switching of the migration mode.

